# Flexing fish: cell surface plasticity in response to shape change in the whole organism

**DOI:** 10.1101/2023.04.28.538646

**Authors:** TE Hall, N Ariotti, HP Lo, C Ferguson, N Martel, Y-W Lim, J Giacomotto, RG Parton

**Affiliations:** Institute for Molecular Bioscience, The University of Queensland, Brisbane, QLD 4072, Australia; Centre for Microscopy and Microanalysis, The University of Queensland, Brisbane, QLD 4072, Australia; Griffith Institute for Drug Discovery, Centre for Cellular Phenomics, School of Environment and Science, Griffith University, Brisbane, QLD 4111, Australia; Queensland Brain Institute, The University of Queensland, Brisbane, QLD 4072, Australia

## Abstract

Plasma membrane rupture can result in catastrophic cell death. The skeletal muscle fibre plasma membrane, the sarcolemma, provides an extreme example of a membrane subject to mechanical stress since these cells specifically evolved to generate contraction and movement. A quantitative model correlating ultrastructural remodelling of surface architecture with tissue changes *in vivo* is required to understand how membrane domains contribute to the shape changes associated with tissue deformation in whole animals. We and others have shown that loss of caveolae, small invaginations of the plasma membrane particularly prevalent in the muscle sarcolemma, renders the plasma membrane more susceptible to rupture during stretch^1–3^. While it is thought that caveolae are able to flatten and be absorbed into the bulk membrane to buffer local membrane expansion, a direct demonstration of this model *in vivo* has been unachievable since it would require measurement of caveolae at the nanoscale combined with detailed whole animal morphometrics under conditions of perturbation. Here, we describe the development and application of the “active trapping model” where embryonic zebrafish are immobilised in a curved state that mimics natural body axis curvature during an escape response. The model is amenable to multiscale, multimodal imaging including high-resolution whole animal three-dimensional quantitative electron microscopy. Using the active trapping model, we demonstrate the essential role of caveolae in maintaining sarcolemmal integrity and quantify the specific contribution of caveolar-derived membrane to surface expansion. We show that caveolae directly contribute to an increase in plasma membrane surface area under physiologically relevant membrane deformation conditions.

## RESULTS

### Development of an *in vivo* system for physiological modulation and analysis of cell and tissue morphometry

We aimed to develop an *in vivo* system where we could induce a defined membrane stretch using a simple physiologically-relevant perturbation, at the same time precisely define changes in morphology at whole animal, tissue, and cellular levels but also quantitate surface domains at the nanoscale. The perturbation should be easily applied to the whole animal, be reproducible and allow comparison to an internal control within the same animal. We identified the zebrafish “active trapping model” as a suitable system fulfilling these criteria.

Under standard conditions, zebrafish embryos begin to hatch from the chorion between two and three days post fertilisation (dpf). However mild bleach treatments, commonly used as a method for pathogen control, have been demonstrated to harden the chorion without impacting zebrafish development. This results in the delayed hatching of a proportion of embryos to 5dpf, with the remainder unable to hatch without manual dechorionation^4^. While trapped within the chorion, embryos display a curvature of the body axis, such that one side undergoes compression while the contralateral side is stretched or extended (Figure 1A). We used this model, which we have termed “active trapping”, as an *in vivo* system to modulate tissue architecture in a specific controlled manner in order to correlate cell shape changes with surface architecture.

**Figure 1:**
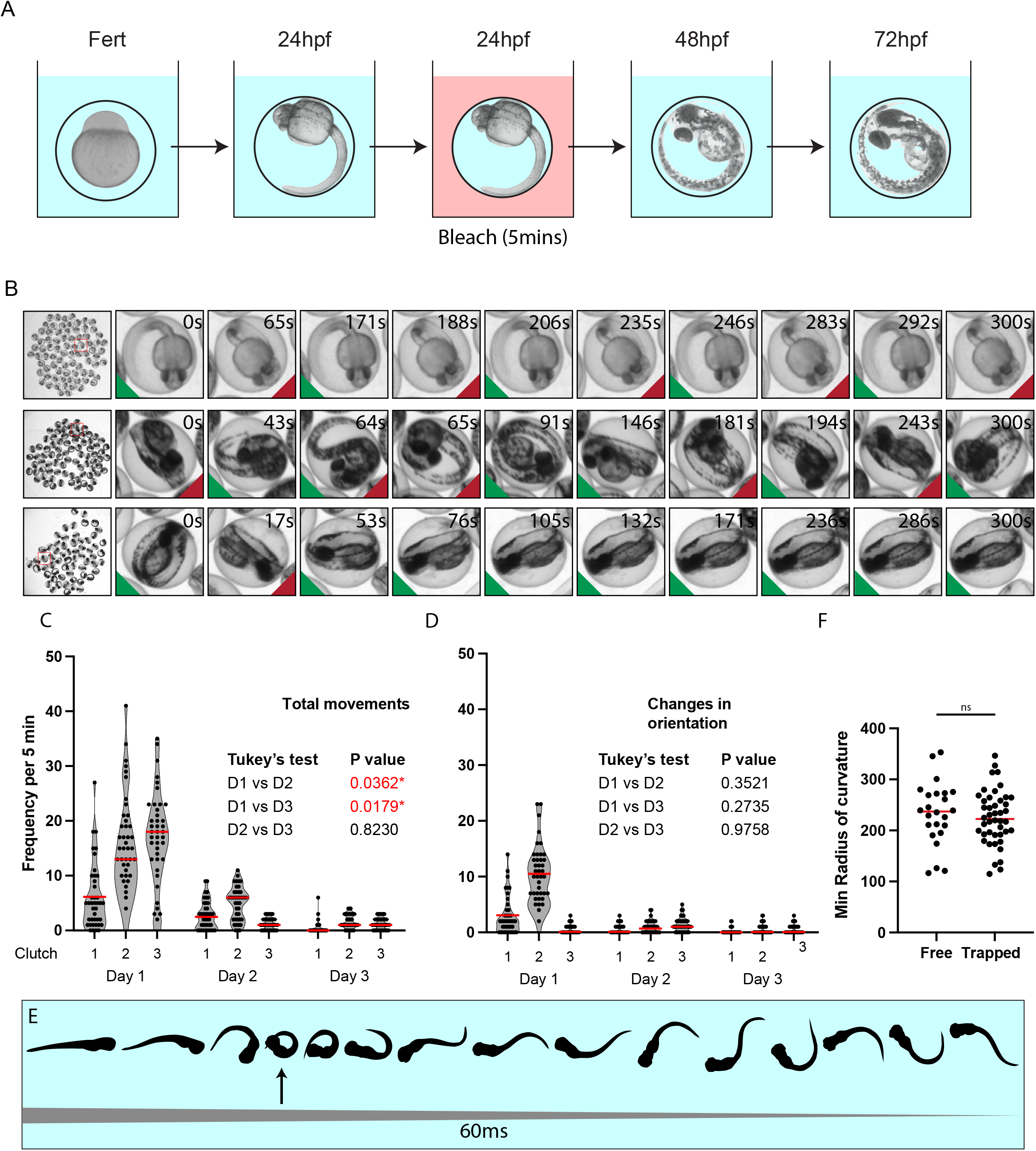
The active trapping system. (A) Embryos are immersed in a dilute bleach solution at 24hpf, which prevents hatching between 48 and 72hpf. Curvature of the body axis is maintained while inside the chorion. (B) Embryos alternate the direction of curvature following spontaneous muscle contractions. Green triangles represent left curvature, red triangles represent right curvature. Time is given in seconds (s). (C & D) Quantitative analysis of total movements and changes in the direction of curvature relative to the body axis. Three clutches were imaged for a period of five minutes on three sequential days beginning at 24hpf (day 1), followed by 48hpf (day 2) and 72 hpf (day 3). n = 40 individuals per clutch. Red bars indicate the mean. Nested one-way ANOVA followed by Tukey’s multiple comparisons test. (E) Silhouettes of a touch-evoked startle response over 60ms showing the point of maximum curvature (arrow). (F) Quantitative analyses of the maximum curvature during fast-starts compared to the maximum curvature generated under active trapping. Two-tailed t-test, n = 24 free, 47 trapped.

We bleached fertilised zebrafish eggs at 24 hours post fertilisation (hpf) and imaged clutches of live embryos at 24, 48 and 72hpf for periods of five minutes. Trapped embryos exhibited sporadic muscle contractions, causing embryos to move within their chorions (Figure 1B & Video S1). Quantitative analyses (Figures 1C & D) showed that these contractions occurred at an average of once every 22s at 24hpf (13.5 times in 5min), resulting in a switch of the direction of curvature with 60% of movements. Contractions declined to once every 84 seconds at 48hrs (3.5/5min, 27% of movements resulting in a change of curvature) and every 279 seconds at 72hrs (1/5min, 41% change in direction of curvature). Embryos could be manually released from their chorions after 72hpf with no deleterious effects on survival (n = 10 clutches, 80 embryos per clutch; % survival = 99.87 at 120hpf/first feeding).

In order to examine the physiological relevance of this process, we imaged touch-evoked startle responses in hatched 3dpf zebrafish using a high-speed camera at 240 frames per second. In response to a sensory stimulus (such as being touched with the tip of a pair of forceps) larval zebrafish exhibit a stereotypical startle response^5^, where the body axis is rapidly contorted into large initial flexion known as the C-bend^6^, followed by subsequent diminishing flexions (Figure 1E, Video S2). We used this footage to measure the maximum curvature (equal to the minimum *radius of curvature*) during the C-bend and found that the curvatures generated by active trapping were remarkably similar to those exhibited during fast-starts at 3dpf (Figure 1F). We conclude that the active trapping protocol is a physiologically relevant system, since the extent of curvature generated occurs both in the eggs of 3-day old unhatched fish and during the larval escape response.

### Light microscopic morphometric analysis of muscle fibres in actively trapped embryos

The active trapping protocol maximised the different morphometry of the trapped embryo at the outward facing surface (convex, extended muscle fibres) and inner surface (concave, compressed muscle fibres). We first sought to quantitatively analyse the morphological changes in skeletal muscle fibres in the compressed and extended sides of the zebrafish embryos. In order to examine the effect of stretch in the live embryo we used a transgenic animal which expresses a membrane targeted GFP in all tissues^7^. In all experiments the same embryo was compared on the extended and compressed sides to reduce inter-animal variability. Optical slices through multiple embryos at 3dpf showed that somite width was on average 20% shorter on the compressed side of the embryo compared to the stretched (Figure 2A, 79 μm vs 99μm). We next used a double transgenic line to measure sarcomere length across multiple sarcomeres using Lifeact-GFP (which marks the thin filament) and the sarcolemma/cell shape by using mCherry-CaaX as a surface marker^8^. These measurements showed that sarcomere length in stretched fibres (2.04μm) was increased by an average of 8% relative to the compressed side (1.89 ± 0.05μm) or to control fibres in resting embryos (1.89 ± 0.03 μm) (Figures 2B & C). This suggested that at any time examined, the muscle fibres were not in an active state of contraction, but that the morphological constraints of the chorion resulted in the outward-facing surface being extended.

**Figure 2:**
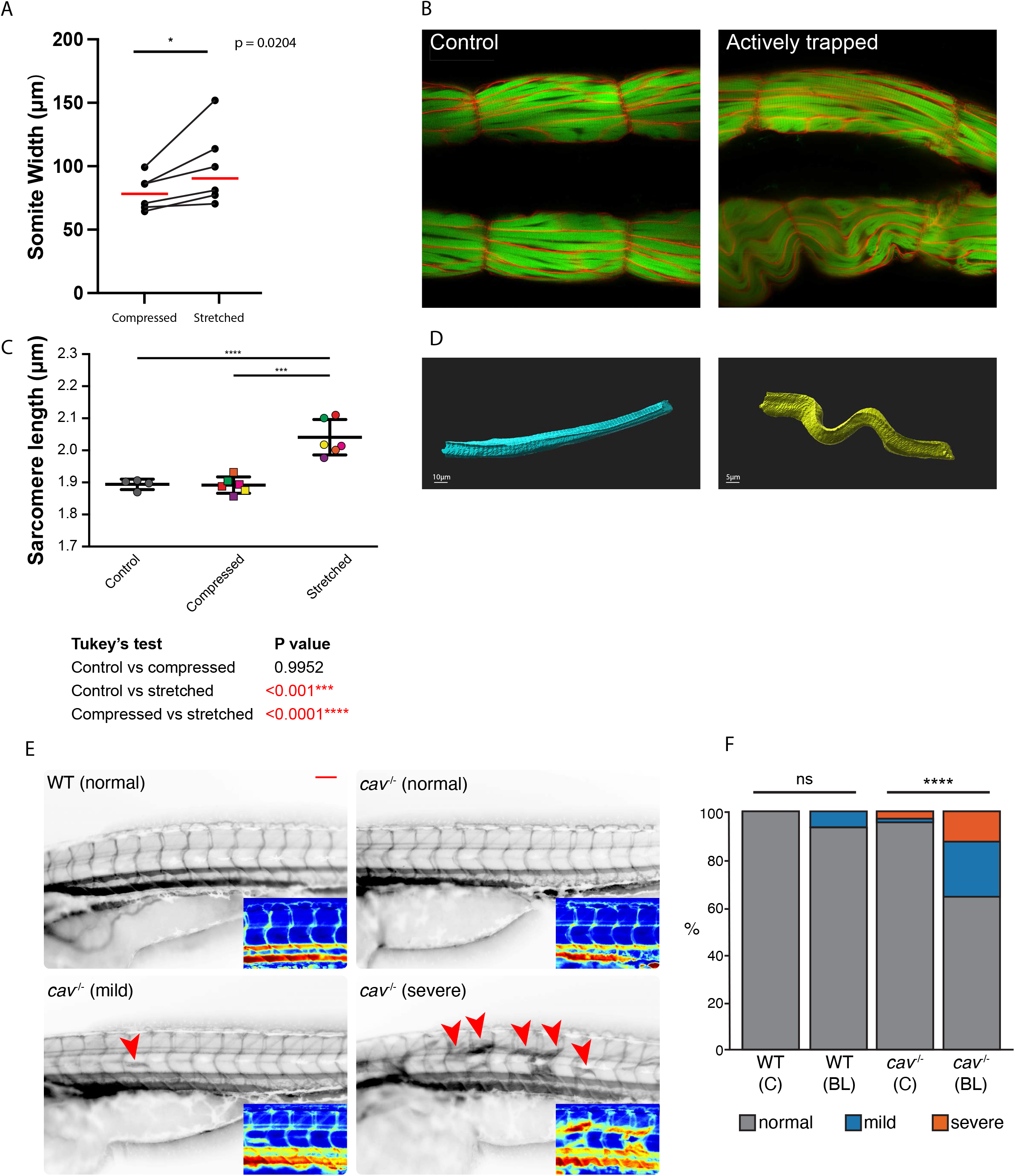
Morphometric analyses in actively trapped embryos. (A) In actively trapped embryos, somite width is increased by 20% on the stretched relative to the compressed side. Red bars indicate the mean. (B & C) Measurements of sarcomere length from actively trapped lifeact-GFP, mCherryCaaX animals show on an increase of 8% on the stretched side relative to at rest. However, sarcomere length is unchanged on the compressed side relative to untrapped controls demonstrating that these fibres are in a relaxed state but not actively contracting. One-way ANOVA followed by Tukey’s multiple comparisons test. Error bars show mean and standard deviation (D) Segmentation of spinning disk images shows that fibres buckle on the compressed side. (E & F) Caveolin loss-of-function models show a higher proportion of Evans Blue infiltration into fibres indicative of membrane failure (arrowheads). Inverted fluorescence images, arrows indicate muscle fibres with uptake. Inset crop with heap map lookup table applied. Fisher’s exact test, *p* < 0.0001. WT = Wildtype C = control (unbleached) BL = bleached. n = 30 WT (C), n = 31 WT (BL), n = 69 *cav*^*-/-*^ (C), n= 56 *cav*^*-/-*^ (BL). Pooled data from three replicate clutches, individual clutches are shown in Figure S2.

A shorter somite width in compressed somites compared to control somites, but no change in sarcomere length, was explained by fibres on the compressed side undergoing buckling as somite width decreased (Figure 2A &B), consistent with the observation that sarcomere length (and therefore total myofibril length) was unchanged from the resting state (Figure 2C). This was confirmed by segmentation of individual fibres from 3-dimensional stacks taken using spinning disk microscopy on GFP-CaaX embryos (Figure 2D). Since only small numbers of fibres were able to be processed in this way, inter-fibre variability using this approach was too great to directly compare differences in cell volume/surface area between stretched and compressed sides. However, fixed sections of individual actively trapped embryos showed a reduction in the minimum Feret diameter of muscle fibres from stretched versus compressed sides of the embryo (Figure S1).

### Genetic ablation of muscle caveolae compromises muscle sarcolemmal integrity in actively trapped embryos

Mutations in *CAV3* result in a loss of muscle caveolae and, in humans, are causative for a muscular dystrophy^9^. In order to model this in our system we first generated a null Cav3 mutant zebrafish. Since *Cav3* is duplicated in zebrafish (into *Cav3* and *CavY* paralogues), we generated *Cav3/CavY* double homozygous knockout embryos.

To assess the physiological consequences of loss of caveolae with respect to muscle cell integrity we compared WT embryos to *Cav3*^*-/-*^*/CavY*^*-/-*^ with active trapping. A membrane impermeable dye, Evans Blue, was injected at 3dpf to evaluate the extent of membrane damage. In this model, intracellular accumulation of EBD indicates impaired membrane integrity^10^. We quantified the number of embryos exhibiting mild or severe EBD uptake according to whether they were wildtype/mutant and/or actively trapped (Figures 2E & F, S2). We found an increase in the number and severity of embryos exhibiting EBD uptake in fibres in the *Cav3*^*-/-*^*/CavY*^*-/-*^ double mutant group, and this was dramatically increased in mutants undergoing active trapping. We conclude that loss of essential caveolar components in *Cav3*^*-/-*^*/CavY*^*-/-*^ mutants predisposes caveolae-deficient cells to plasma membrane rupture during mechanically-induced cell shape change, validating the use of the active trapping model for studies of mechanical compensation *in vivo*.

### A predictive model of membrane dynamics during muscle fibre stretch

In order to generate a predictive model of membrane stretch in muscle fibres under active trapping, we began by defining a model population. We generated a three-dimensional confocal volume from a hatched wildtype embryo at 3df and used this to define the muscle fibre cellularity profile (distribution of cross-sectional areas). To measure from the centre of the fibres the volume was optically resliced at 45 degrees through the centre of the myotome, then rescaled back to the correct aspect ratio (Figure S3A). Fibre morphometrics were measured from successive epaxial somites using our custom written macros as previously described^11^. The defined distribution of cross-sectional areas was then used to generate an idealised myotomal map, using a custom script, which populates a model myotome with cells in a spatially stochastic pattern with a bias towards smaller diameter cells at the periphery. Cell boundaries are then generated using Voronoi transformation^12^. Using this model myotome we were then able to examine the effect of applying differential stretch *in silico* to all fibres within the myotome, with the assumption that the volume of a fibre does not change when compressed or stretched, since liquid is incompressible (Figure 3A). According to our model, in an idealised population fibres would be stretched by between 0.5-18% depending on position within the myotome, with an average of 11%. This is in close agreement with our empirically measured data on sarcomere length (Figure 2C). This stretch would generate an average increase of 4.6% in surface area (max. 8.4%, min. 0.3%). On the contralateral side under compression, the total surface area of a fibre would not change since fibres buckle rather than shorten, although we would predict greater local heterogeneity between different regions of the plasma membrane of each cell.

**Figure 3:**
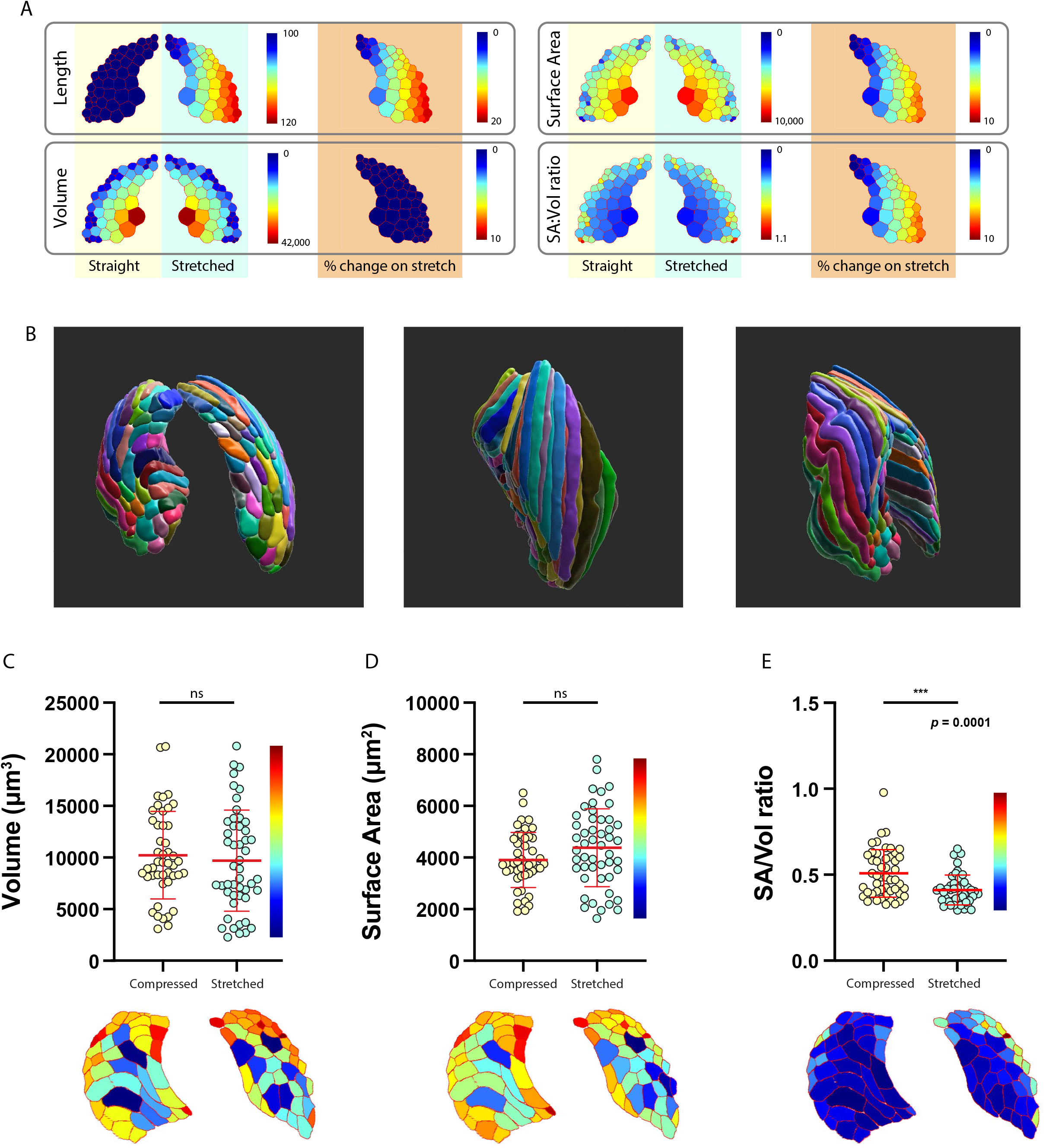
A predictive model of fibre stretch. (A) *In silico* prediction of fibre morphometrics after the applicaton of stretch. (B) Three dimensional rendering of a (real) myotome under stretch generated by hand segmentation in order to validate the modelling approach. In this model fibres are colour coded randomly as a visual aid and do not represent any morphometric parameter. See also video S3. (C-E) Empirical measurements of volume (C), surface area (D) and SA:Volume ratio (E) from the dataset shown in (B). Myotomal heatmaps display a virtual transverse section through the rendering and are colour coded according to the data in the graphs. Statistics were two-tailed unpaired t-tests. Error bars indicate the mean and standard deviation.

In order to validate our model, we chose to collect data from an entire epaxial somite (both stretched and compressed sides) in 3-dimensions from a single actively trapped embryo. We used manual segmentation in ImageJ using custom macros to generate rendered volumes for each individual fibre and capture morphological parameters. Volumes were then imported into Imaris software for 3D viewing (Figure 3A & B, S3A & B, Video S3). In agreement with our model, the linear distance between the ends of the fibres, the length along the centroid and the linearity index was significantly greater in stretched versus compressed fibres (Figures S3C-E). The buckling effect can clearly be seen in the 3D rendered volume (Figure 3B). We found that fibre volume was 5% lower on the stretched side versus the compressed side while surface area was increased by 12% (Figures 3C & D). As predicted from our model data, due to the morphologically heterogenous population and small effect size, neither was statistically significant. However, the surface area/volume ratio was significantly lower on the stretched vs compressed side (Figure 3E). Overall, these empirical data support the conclusions from the modelling.

### Caveolar disassembly directly supplies demand for increased membrane area upon cellular stretch

This system offered a unique opportunity to gain quantitative ultrastructural insights into morphological differences in the same muscle fibre types, but in different areas of the same embryo, avoiding any variability between individuals. We could therefore quantitatively compare the ultrastructure of genetically identical muscle fibres exposed to different mechanical forces and directly examine the number of caveolae in different muscle fibres to test our predictions from light microscopy. We prepared actively-trapped embryos and processed them for conventional transmission electron microscopy (TEM) and serial blockface scanning electron microscopy (SBF-SEM). Thin section TEM confirmed a greater mean sarcomere length between concave and convex sides of individual embryos (concave/compressed side sarcomere length of 1.93 ± 0.23, extended/convex side sarcomere length 2.03 ± 0.06μm; not shown).

We next prepared intact trapped embryos for whole embryo sectioning. Embedded embryos were mounted within the chamber of the SBF-SEM so that the embryo was sequentially imaged from the convex side of the fish, through the centre of the embryo (notochord) and then through to the concave side (over 3000 sequential images, 100nm slice thickness; Figure 4A, S4A, Video S4). With the aim of examining surface caveolae, we defined conditions that would allow recognition and quantification of all caveolae within a 3dpf muscle fibre *in situ*. For these analyses we defined caveolae as surface-connected pits of ∼100nm diameter (Figure S4B, Video S5).

**Figure 4:**
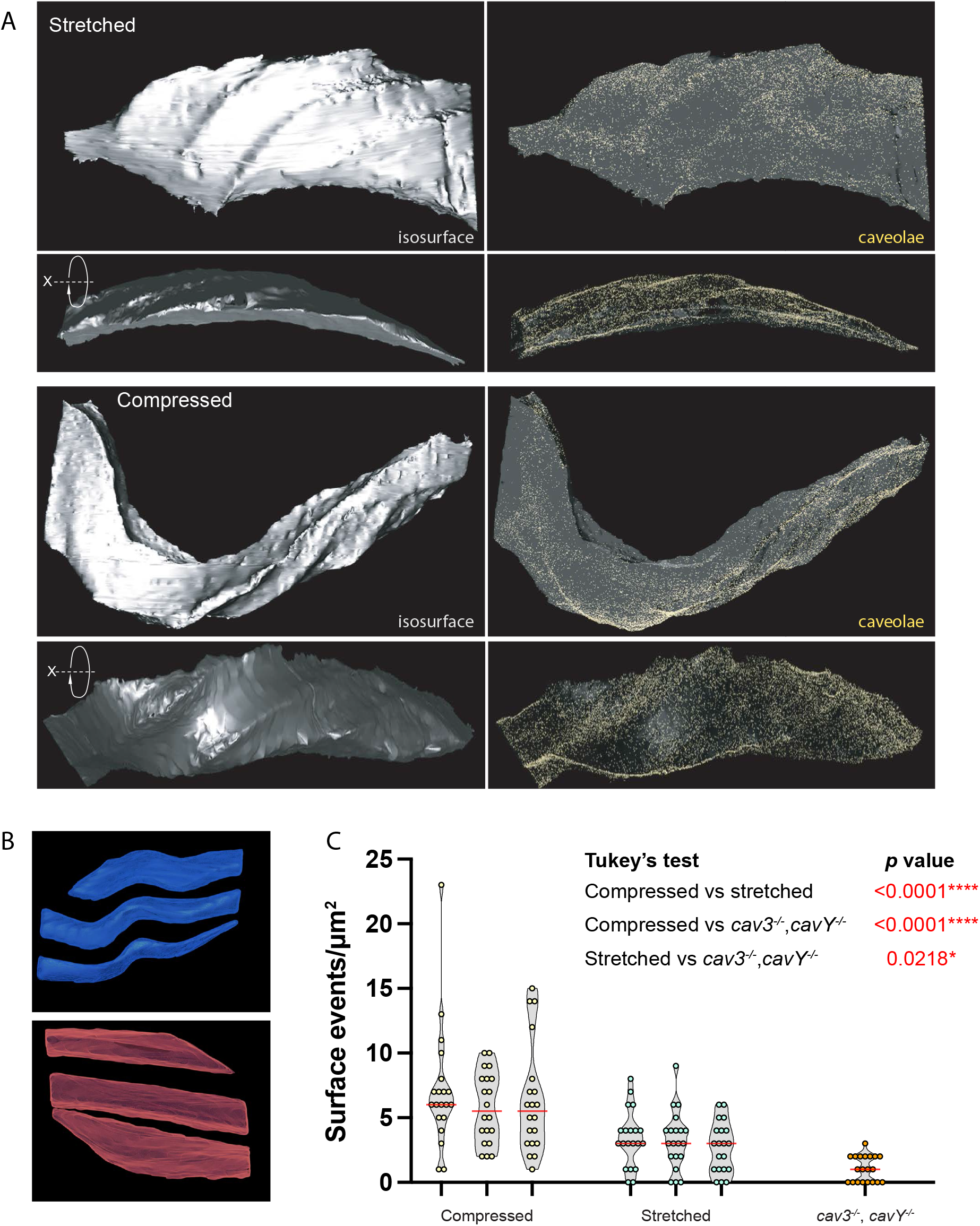
Quantitative analysis of caveola number under active trapping. (A) Rendering of caveolae in 3d dataset in stretched vs compressed fibres. (B) Three dimensional rendering of multiple stretched (red) vs compressed (blue) fibres from the same serial blockface electron microscopy dataset. (C) Quantification of surface caveolae from stretched and compressed fibres, and caveolae-deficient *cav3*^*-/-*^,*cavY*^*-/-*^ mutants. Red line indicates the mean. Statistics were a nested one-way ANOVA with Tukey’s multiple comparison test.

We performed quantitation of caveolae from representative compressed and stretched fibres (Figures 4B & C). Stretched fibres showed a striking reduction in caveolar density as compared to compressed fibres within the same embryos; an individual 3dpf zebrafish myofibre has a caveolar density of 6.42 caveolae/μm^2^ on the compressed side compared to 3.12 caveolae/μm^2^ on the stretched side (note also the dramatic reduction in surface connected caveola-like structures in *cav3*^*-/-*^/*cavY*^*-/-*^ null animals, 3.12μm^2^ stretched, 6.42/μm^2^ compressed, 1.05/μm^2^). In our model fibre population we calculated that the average initial resting dimensions of 100μm in length with an average surface area of 4541μm^2^, under active trapping, the average surface area would increase by 200μm or 4.4%. According to our measurements of caveolar density in stretched vs compressed lateral fibre membranes (3.12μm^2^ vs 6.42μm^2^), and assuming a single caveola has a surface area of 0.0114μm^2^ (calculated using a 60nm diameter vesicular structure), the observed reduction in caveolae would provide an additional 176μm^2^ of surface membrane (+3.73%).

## DISCUSSION

In this study we have developed a physiologically relevant system in which it is possible to induce mechanical deformation of tissues and cells within a live organism and conduct quantitative multiscale analyses in a wholistic manner. We have used this system to directly examine the effect of membrane stretch on caveolar disassembly, but these approaches could be equally effective for other studies of the effects of mechanical changes on cellular function, requiring a simultaneous analysis of *in vivo* architecture at the macroscale, cell morphology at the microscale and ultrastructural architecture at the nanoscale. Previously, we and others have demonstrated a reduction in caveola density in cultured cells (including muscle fibres), in response to changes in membrane tension^1–3,13^ and there is evidence for reduced caveolae numbers in cardiomyocytes and endothelial cells following increased cardiac output^3^. In addition, loss of caveolar components *in vivo* leads to membrane failure in zebrafish skeletal muscle^13^ and the notochord^4,14^ and human pathogenic variants in caveolar proteins result in muscular^15^ and lipo-dystrophies^16,17^. The active trapping model allows us to move from the whole organism under curvature, through individual segmented cells, to caveolae in a quantitative manner. Using a combination of transgenic zebrafish under active trapping with fluorescence-based morphological assays and high resolution quantitative electron microscopy we have built a uniquely detailed picture of the role of caveolae in cell shape plasticity. Our empirical data and modelling show that active trapping results in an almost instantaneous increase in sarcolemma surface area on the extended side of the zebrafish. This is coupled with an approximately two-fold reduction in caveolar density. Strikingly, we find that this increase in surface area closely matches the amount of membrane provided by the loss of caveolae from the sarcolemma, demonstrating that almost all additional membrane required during muscle fibre stretch *in vivo* is contributed directly by caveolae.

## STAR*METHODS

### Zebrafish maintenance

Zebrafish embryos were sourced from The University of Queensland Biological Resources (UQBR) breeding colony where they were maintained according to standard conditions^18^ under institutional guidelines approved by the University of Queensland (UQ) Animal Ethics committee (Tecniplast recirculating system, 14-h light/10-h dark cycle). Adults (90 dpf above) were housed in 3 or 8 L tanks with flow at 28.5°C, late-larval to juvenile stage zebrafish (6 dpf to 45 dpf) were housed in 1 L tanks with flow at 28.5°C and embryos (up to 5 dpf) were housed in 8 cm Petri dishes in standard E3 media (5 mM NaCl, 0.17 mM KCl, 0.33 mM CaCl_2_, 0.33 mM MgSO_4_) at 28.5°C (incubated in the dark). The following zebrafish strains were used in this study: wild-type (AB), three transgenic strains were used in this study; a ubiquitously expressed membrane GFP (*actb2:EGFP-CaaX*^*pc10tg*^)^7^, ubiquitously expressed Lifeact-GFP (*actc1b:Lifeact-EGFP*^*pc21tg*^)^8^ and a muscle specific membrane mCherry (*actc1b:mCherry-CaaX*^*pc22tg*^)^8^. The developmental stages of zebrafish used in experiments (up to 5 dpf) are prior to specific sex determination^19^ and are specifically stated in figures. All zebrafish used in experiments were healthy, not involved in previous procedures and were drug and test naive.

The zebrafish *Cav3*^*uq12rp*^ mutant line (g.15C>A; p. Y5X) was generated by screening an ENU mutagenesis library at the Hubrecht Institute, Utrecht, Netherlands. The *CavY*^*uq13rp*^ mutant line (g.46delG; p.L55X) was generated using Golden Gate TALEN technology^20^ targeting the following sequence: TCCAGTGAGACTGAGATCGAcctgagggacgcagGAGACGGAGAGGACGACGA.

### Chorion bleaching

Chorion bleaching was carried out by immersing 24hpf embryos in 0.006% bleach (30μl 10% hypochlorite per 50mls of E3 media) for 5 minutes^4^. Embryos were subsequently washed three times in fresh E3 which was changed daily.

### Recording of flips/twitches

In qualifying total movements and the number of changes in orientation, timelapse photomicroscopy was carried out via brightfield imaging using a Nikon SMZ1500 fluorescence stereomicroscope in groups of 80 embryos per dish. 1880 frames were captured over 5 minutes (frame rate 6.3 per second). Movements and changes in orientation were recorded manually on playback.

### Measuring radius of curvature

Radius of curvature was measured using Kappa plugin in ImageJ.

### Touch evoked fast start assay

Single embryos were placed in dishes filled with E3 media. Tactile stimuli were applied to the head of the embryo with a pair of no. 5 forceps to elicit a response. Footage was recorded using a with a Sony A7S III camera in XAVC HD 50M 4:2:0 8bit format with a shutter speed of 0.0005 seconds at 240 frames per second. Movies were exported at 30 frames per second and spatial calibration was achieved using an in frame S81K Stage micrometer.

### Vibratome sections

Vibratome sectioning was carried out as previously described^21^. Zebrafish embryos were fixed in 4% PFA overnight at 4°C and mounted in 8% low-melting-point agarose (#A9414; Sigma-Aldrich). Vibratome sections were cut to a thickness of 150 μm using a Leica VT1000S vibratome and mounted in Mowiol (#475904; Merck). Confocal images were captured on a Zeiss Axiovert 200 upright microscope stand with LSM710 meta confocal scanner (63× Plan Apochromat objective NA 1.40).

### Live imaging of zebrafish embryos

Live imaging was carried out as previously described^22^. Zebrafish embryos were anesthetized in tricaine (#A5040; Sigma-Aldrich)/E3 solution and mounted in 1% low-melting-point agarose (#A9414; Sigma-Aldrich). Confocal images of mounted embryos (immersed in tricaine/E3) were captured on three different microscopes; an upright Zeiss LSM710 meta confocal (40× Plan Apochromat water immersion objective NA 1.0), an upright Zeiss LSM900 confocal (40× Plan Apochromat water immersion objective NA 1.0) and an Andor Dragonfly spinning disk confocal (Apo lamda LS 40× immersion objective NA 1.15).

### Image Processing and quantification

All image processing was carried out in Fiji (ImageJ)^23^ and figures were prepared in Adobe Illustrator 2023. A number of custom written ImageJ macros were used in the analyses and full code will be deposited in Zenodo with stable DOIs upon publication.

### Evans Blue Dye injections and scoring

Evans blue injections were carried out at 72-hpf by injection of approximately 5nL of 0.1mg/ml into the pre-cardiac sinus^24^. Embryos were incubated at 28°C for four hours to ensure sufficient circulation an uptake before dechorionation and imaging. Embryos were classified as normal, mild or severe on the basis of Evans blue uptake into muscle fibres.

### Serial Block face Scanning Electron Microscopy

3dpf zebrafish embryos were anaesthetised with tricaine and initially fixed in 2.5% glutaraldehyde in a BioWave microwave (Pelco). Zebrafish were then dechorionated and refixed in 2.5% glutaraldehyde prior to processing for serial block face scanning electron microscopy as performed previously (Hall et al, 2020). Briefly, zebrafish were postfixed in 2% Osmium Tetroxide (OsO_4_) with 1.5% Potassium ferricyanide. Samples were washed in water, incubated in 1% thiocarbohydrazide for 20 minutes, postfixed in 2% OsO_4_ in the BioWave, washed, stained *en bloc* with 1% uranyl acetate and subsequently with lead aspartate (20mM lead nitrate, 30mM aspartic acid, pH 5.5). Embryos were serially dehydrated in acetone, infiltrated with durcopan resin and polymerised. Blocks were mounted for acquisition on a Zeiss Gemini FE-SEM fitted with a Gatan 3view, with a magnification of 7.5K, pixel size of 5.7nm, at high vacuum. Sections were 100nm in thickness.

### Segmentation and quantification of 3D EM data

Serial block face scanning electron micrographs were processed using the IMOD suite of image processing tools^25^. Stacks were aligned using “Align serial sections” in Etomo and surface area measurements were extracted using Imodinfo. Caveolar quantifications were performed as follows; three cells and 20 regions per cell were selected from the extended and compressed sides. Each region comprised ∼2 μm^2^ in cross-sectional area from similar regions of the sarcolemma. Caveolae were defined as surface connected vesicular structures greater than 50nm but less than 200nm in diameter. Identical quantification was performed on the Cav3/CavY control zebrafish to give a baseline reading of surface connected pit like structures in a caveolae null background.

### Statistics

All statistics were carried out in Prism9 for macOS. Specific tests used and exact *p* values are given in the figure legends. Where *p* was less than 0.0001, “*p* < 0.0001” is stated.

## ACKNOWLEDGEMENTS

We are grateful to Jeroen Bakkers (Hubrecht Institute, Utrecht, Netherlands) and Susan Nixon (Institute for Molecular Bioscience, Brisbane, Australia) for generation of the *Cav3*^*uq12rp*^ mutant zebrafish line, and to Neil Bower (Institute for Molecular Bioscience, Brisbane, Australia) for design and generation of the TALEN vectors used in production of the *CavY*^*uq13rp*^ mutant line. This work was supported by the National Health and Medical Research Council of Australia (grant APP1099251 to T.E.H. and R.G.P. and fellowship APP1156489 to R.G.P.) and the Australian Research Council (grant DP200102559 to T.E.H. and R.G.P). The authors acknowledge the use of the Microscopy Australia Research Facility at the Centre for Microscopy and Microanalysis at The University of Queensland. Confocal microscopy was performed at the Australian Cancer Research Foundation (ACRF) and the Institute for Molecular Bioscience (IMB) Dynamic Imaging Facility for Cancer Biology with funding from the ACRF.

## AUTHOR CONTRIBUTIONS

Conceptualisation, RGP, TEH, NA Data curation RGP, TEH, NA, HL, CF, GJ Formal analysis RGP, TEH, NA, HL, CF, GJ Funding acquisition RGP, TEH Investigation RGP, TEH, NA, HL, GJ, NM, Y-W L Methodology RGP, TEH, NA, HL, CF, NM, Y-W L Supervision RGP, TEH Visualisation RGP, TEH, HL Writing – original draft RGP, TEH Writing – review and editing RGP, TEH, NA, HL

---

**Figure S1:**
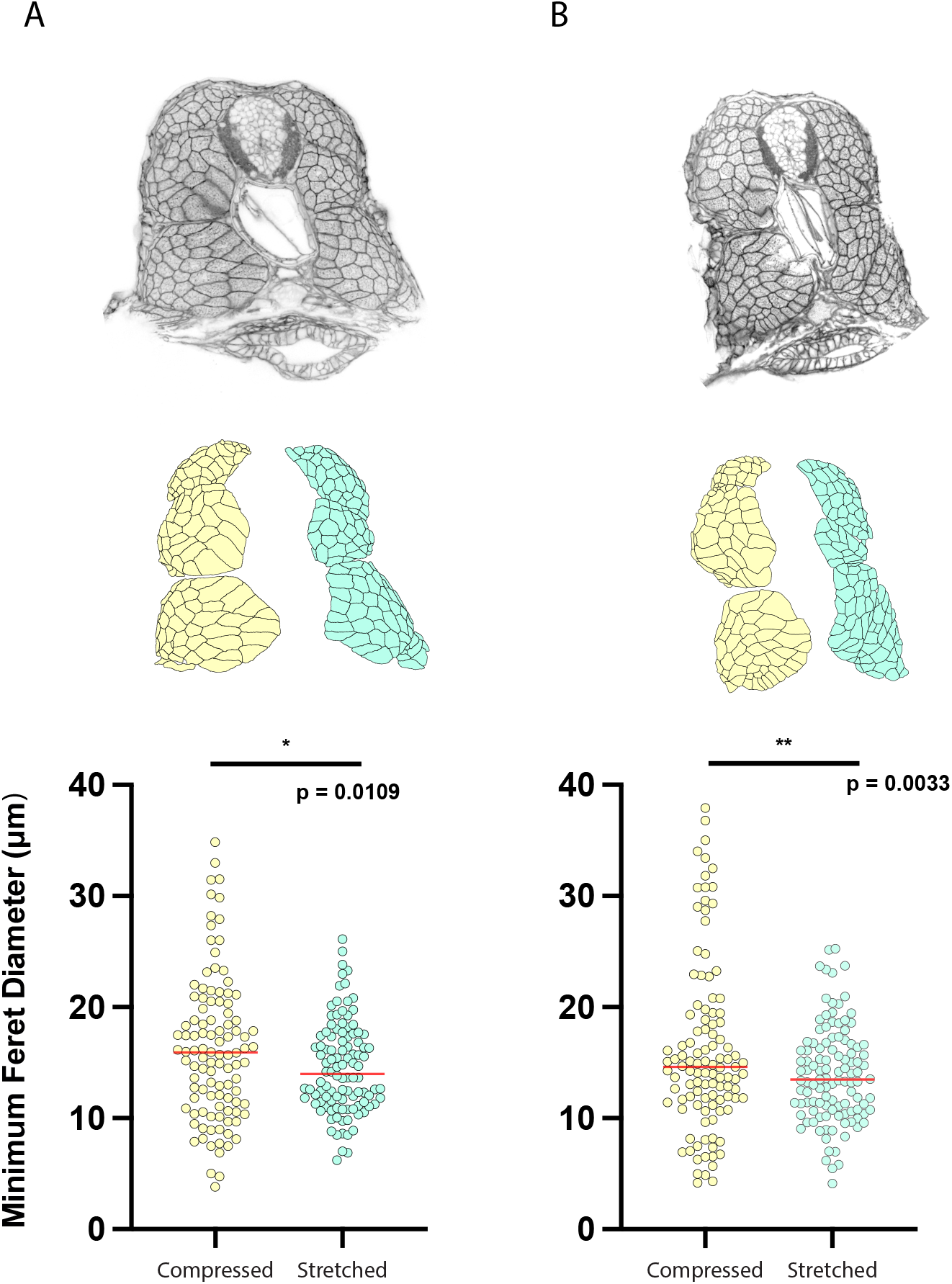
Cellularity analysis of actively trapped embryos using vibratome sections. The minimum Feret diameter of muscle fibres is between 10 and 15% greater on the compressed side relative to the stretched side. Statistics were one-way ANOVA followed by Tukey’s multiple comparisons test. A and B show individual embryos. Red bars indicate the mean.

**Figure S2:**
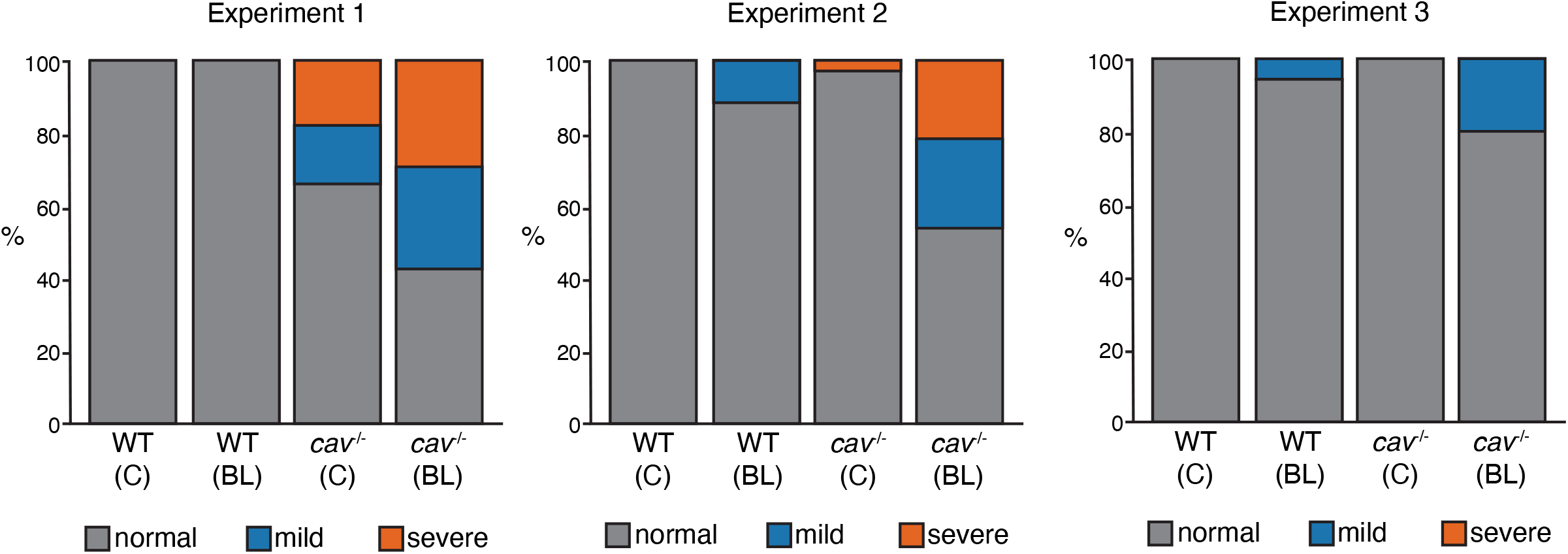
A higher proportion of *cav3*^*-/-*^,*cavY*^*-/-*^ mutant embryos show uptake of Evans blue dye (EBD) in response to chorion bleaching. Individual clutch data from Figure 2F.

**Figure S3:**
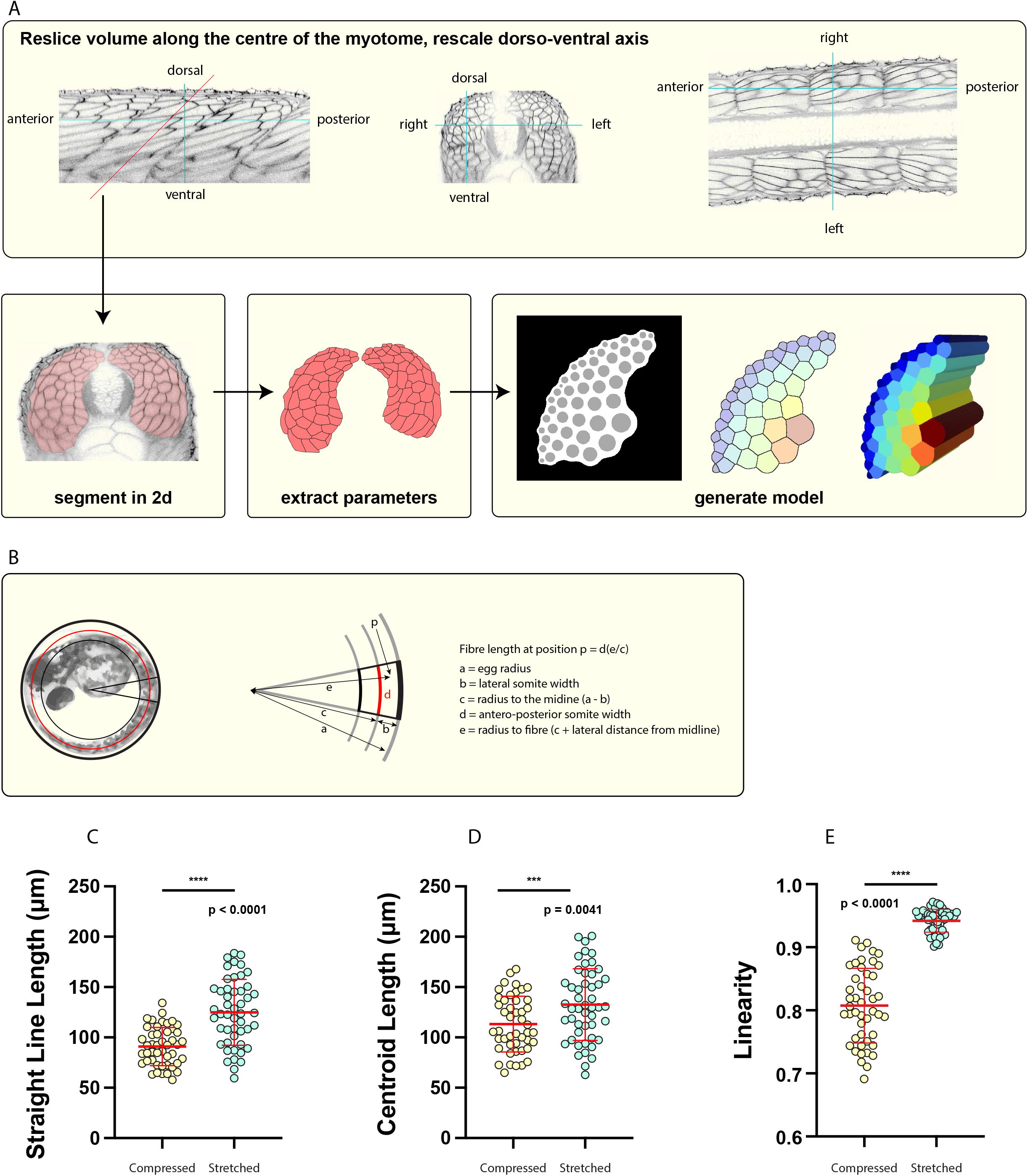
The effect of stretch on muscle fibres under active trapping. (A) In order to define the cellularity profile from the centre of each fibre, a three-dimensional confocal volume was computationally resliced at 45 degrees and rescaled back to the correct aspect ratio. Fibres were then traced by hand in ImageJ. Model three dimensional myotomes were then generated from the cellularity profile using a custom written script that places fibres semi-randomly in the myotome according to size (with the smallest fibres occupying the outside layer). Fibre boundaries are placed using a Voronoi transformation. (B) Stretch is modelled as a function of four variables, egg radius, lateral somite width, antero-posterior somite width, and the lateral distance of the fibre from the midline. (C-E) Empirical measurements of the linear distance between fibre termini from the dataset shown in Fig. 3B. (C) the straught line distance between fibre termini. (D) distance between fibre termini measured down the centroid of the fibre. (E) fibre linearity (straight line length/centroid length). Statistics were two-tailed unpaired t-tests. Error bars indicate the mean and standard deviation.

**Figure S4:**
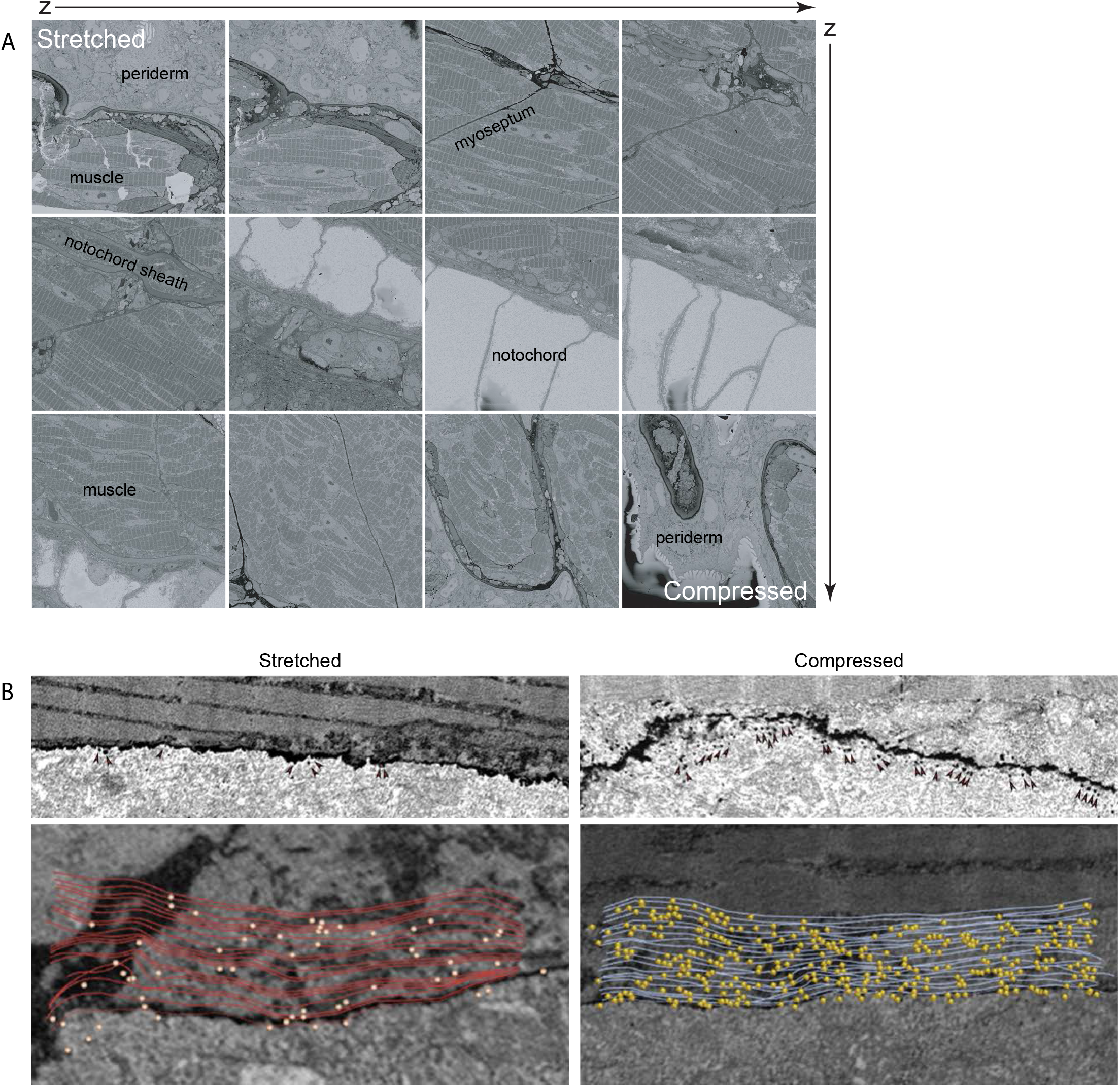
Modelling caveolar dissassembly at the ultrastructural level. (A) Stills from the serial blockface electron microscopy dataset of an actively trapped embryo, moving from the periderm on one side, through the entire trunk to the periderm on the other side. See also video S4 (B) Representative areas from the 3d dataset in stretched vs compressed fibres. Top panels - caveolae are indicated by arrowheads. Bottom panels - micrographs overlaid with segmented caveolae (yellow) and orientation of overlying membrane (lines).

